# Dynamic dot displays reveal material motion network in the human brain

**DOI:** 10.1101/2020.03.09.983593

**Authors:** Alexandra C. Schmid, Huseyin Boyaci, Katja Doerschner

## Abstract

There is growing research interest in the neural mechanisms underlying the recognition of material categories and properties. This research field, however, is relatively more recent and limited compared to investigations of the neural mechanisms underlying object and scene category recognition. Motion is particularly important for the perception of non-rigid materials, but the neural basis of non-rigid material motion remains unexplored. Using fMRI, we investigated which brain regions respond preferentially to material motion versus other types of motion. We introduce a new database of stimuli – dynamic dot materials – that are animations of moving dots that induce vivid percepts of various materials in motion, e.g. flapping cloth, liquid waves, wobbling jelly. Control stimuli were scrambled versions of these same animations and rigid three-dimensional rotating dots. Results showed that isolating material motion properties with dynamic dots (in contrast with other kinds of motion) activates a network of cortical regions in both ventral and dorsal visual pathways, including areas normally associated with the processing of surface properties and shape, and extending to somatosensory and premotor cortices. We suggest that such a widespread preference for material motion is due to strong associations between stimulus properties. For example viewing dots moving in a specific pattern not only elicits percepts of material motion; one perceives a flexible, non-rigid shape, identifies the object as a cloth flapping in the wind, infers the object’s weight under gravity, and anticipates how it would feel to reach out and touch the material. These results are a first important step in mapping out the cortical architecture and dynamics in material-related motion processing.

## 1. INTRODUCTION

Recognizing and estimating the material qualities of objects is an essential part of our visual experience. Perceiving material qualities quickly and correctly is critical for guiding decisions and actions, whether we are deciding what fruit to eat, if a blanket is soft enough, or how we should grip a porcelain cup. Despite the importance of recognizing and estimating the properties of materials, it is still not well understood how the brain accomplishes these tasks and, compared to object and scene perception, neuroscientific studies investigating material perception are relatively recent (see Komatsu & Goda, 2018 for a review).

Literature investigating the neural mechanisms of visual material perception has so far heavily focused on how the brain discriminates between different surface optical appearances; that is, the cortical areas involved in the visual processing of the surface properties of materials (Komatsu & Goda, 2018). These properties, such as a surface’s micro- and meso-structure, or its reflective-, transmissive- and refractive properties, give a material its characteristic visual appearance (e.g. glossy, plastic-y, metallic, etc.). Nearly all these studies have used static images as stimuli (but see Okazawa et al. 2012; Kam et al. 2015; see Sun et al. 2016a for binocular stimuli).

The use of static images when investigating the visual processing of materials is somewhat justified because, although material perception is a multisensory (Baumgartner et al. 2013), dynamic experience (Schmid & Doerschner, 2019), many material qualities can be conveyed through images alone (Paulun et al. 2017; Schmid & Doerschner, 2018; Schmidt et al. 2017; van Assen & Fleming 2016, Baumgartner et al. 2013, Xiao et al. 2016). But not all material properties can be faithfully conveyed through static images, and most mechanical material properties become much more vividly apparent with motion: watching a rubber band stretching, a jelly jittering, and hair bending elicits strong impressions of elasticity, wobbliness, and softness, respectively. In fact, image motion has been shown to provide information about material qualities over and above the information available from 2D images (Doerschner et al. 2011, Schmid & Doerschner 2018; Schmidt et al. 2017). Furthermore, recognizing materials in natural environments likely entails integrating spatiotemporally segregated information into coherent percepts, similar to detecting animate objects (think of detecting a tiger through long, swaying grass), making it a special kind of event perception (Johansson, 1974; Jannson, Bergström & Epstein, 1994; Radvansky & Zacks, 2011; Bingham, Schmidt & Rosenblum, 1995). Given that many brain regions are sensitive to certain kinds of motion structure (for reviews see Erlikhman et al., 2018; Kourtzi et al., 2008; Nishida et al., 2018), it is surprising that, to date, no studies have examined the cortical processing of non-rigid material motion. One possible reason for this is that it is difficult to find adequate control conditions for complex, dynamic stimuli. Recent improvements in computing power now puts us in a position to be able to simulate complex material motion stimuli in a controlled way and generate adequate control conditions (described below).

The first step to investigate the neural correlates of non-rigid material motion is to find whether cortical areas exist that show a preference for non-rigid material motion over other types of motion. In this study, we developed a new class of stimuli that utilizes the fact that mechanical and tactile qualities of materials can be convincingly conveyed through image motion alone (Schmid & Doerschner, 2018; Bi et al., 2019). Previously these have been called point-light stimuli by Schmid & Doerschner (2018), analogous to “point light walkers” that have been used extensively in biological motion research (Johansson, 1973). Similar stimuli have also been called “dynamic dot stimuli” (Bi et al., 2019). Here, we name our new movie database “**dynamic dot materials**”, where specific non-rigid material types are solely depicted through the motion of black dots on a gray background. Dynamic dot materials not only depict mechanical material properties convincingly; they also isolate dynamic properties from optical (surface) cues and can provide more stimulus control compared to “full cue” videos that contain surface information. This is because the behavior of trajectories and speed of individual dots can be manipulated.

Using fMRI, we investigated which brain regions respond preferentially to dynamic dot materials versus motion-scrambled and structure-from-motion control stimuli. Comparing material motion to these two control conditions allowed us to identify regions that responded to material motion over and above just any motion (scrambled motion condition), and over and above general structure from motion, motion coherence, and bounded “object-ness” (structure-from-motion control stimuli). Anticipating our results, we find widespread preferential activation for non-rigid material motion versus the control conditions. This suggests that dynamic dot materials are very suitable for mapping the cortical network involved in the perception of non-rigid materials, and we recommend that our stimuli can be used in future studies – for example as a localizer – to further explore the specific function of these different regions in this “material motion network”.

## 2. MATERIALS & METHODS

### 2.1. Participants

Ten volunteers participated in the experiment (age range: 21-42, 2 males, mean age: 27.3, 1 left handed). All participants had normal or corrected to normal vision and had no known neurological disorders. Participants gave their written consent prior to the MR imaging session. Protocols and procedures were approved by the local ethics committee of Justus Liebig University Giessen and in accordance with the Code of Ethics of the World Medical Association (Deklaration of Helsinki).

### 2.2. Stimuli

#### 2.2.1. Dynamic dot materials

Stimuli were non-rigid materials, generated with Blender (version 2.7) and Matlab (release 2012a; MathWorks, Natick, MA) and presented using Matlab and the Psychophysics Toolbox (Brainard, 1997). Each dynamic dot material was a 2s animation (48 frames) that depicted the deformation of a specific material under force. Object deformations were simulated using the Particles System physics engine in Blender, with either the fluid dynamics or Molecular addon (for technical specifications we refer the reader to Schmid & Doerschner, 2018). For variety, we simulated non-rigid materials with different mechanical properties under various forces (details shown in Table 1), including cloth flapping in the wind, liquids rippling and waving, breaking materials of high, medium, and low elasticity, as well as non-breaking elastic materials. Upon creation of these different material types we exported the 3D coordinates of each particle at each frame. Using Matlab, we calculated the 2D projection for each particle from a specific viewing angle. For each material only 200 random particles were selected for display, as a medium density yielded best perceptual impressions in previous work (Schmid & Doerschner, 2018), with the remaining particles set to invisible. The particles were sampled from throughout the volume of the substance with the following exception: for liquids the particles were sampled from the surface because sampling from the volume in these cases did not convey the material qualities as convincingly. In total we rendered ten animations (Table 1). Frames from example stimuli can be seen in Figure 1a, and the trajectory of the dots over time can be seen in Figure 2 (left column) and Supplementary Figure 1. We also informally confirmed with participants that they perceived the dynamic dot materials as intended. We formally recorded four participants’ responses, which are shown in Supplementary Table 1.

**Table 1:** Simulation and rendering details for each material motion animation. Animations were 2s clips of materials being deformed in various ways. The first column shows the name of the movie provided in the Supplementary Material. The second column shows whether each material was simulated as linked particles (breaking or unbreaking) or fluid particles (see main text). The third column shows details of how the simulation was set up in Blender. The fourth column shows what part of the simulation was rendered in the final animation.

| Material motion name | Particle physics | Blender simulation details | Part of simulation rendered |
| --- | --- | --- | --- |
| cloth | Linked, unbreaking particles | A sheet of linked particles was attached at the top of the scene and blown by a wind force field. | Cloth blowing in the wind. |
| Cloth_rot | Linked, unbreaking particles | Same as cloth. | Cloth movie rotated 90 degrees. |
| Ripple | Fluid particles | Fluid particles were dropped into a small round container, causing it to ripple. | Liquid rippling. |
| Waves | Fluid particles | Fluid particles were dropped into a large square container and stirred by an invisible rod causing larger waves. | Liquid waves. |
| pokeWobble | Linked, unbreaking particles | Elastic cube made of linked particles was attached to the ground and poked with an invisible rod. | Cube wobbling. |
| pokeWobble_rot | Linked, unbreaking particles | Same as pokeWobble. | Cube wobbling, rendered from a different camera angle to pokeWobble. |
| stretchBounce | Linked, unbreaking particles | Elastic cube made of linked particles was attached to invisible solid walls that moved horizontally apart. Elastic cube stretched and bounced. | Cube stretching and bouncing. |
| stretchWobble | Linked, breaking particles | Same as stretchBounce but elastic cube ripped and wobbled like hard jelly. | Cube stretching, ripping, and wobbling. |
| stretchHighWobble | Linked, breaking particles | Same as stretchBounce but elastic cube ripped and wobbled substantially. | Cube stretching, ripping, and wobbling. |
| stretchDough_rot | Linked, breaking particles | Same as stretchBounce but the cube was made of low-elastic material that ripped apart softly with no wobble motion. | Cube stretching and ripping. Movie was rotated 90 degrees to provide variation to the other three “stretch” movies. |

**Figure 1.**
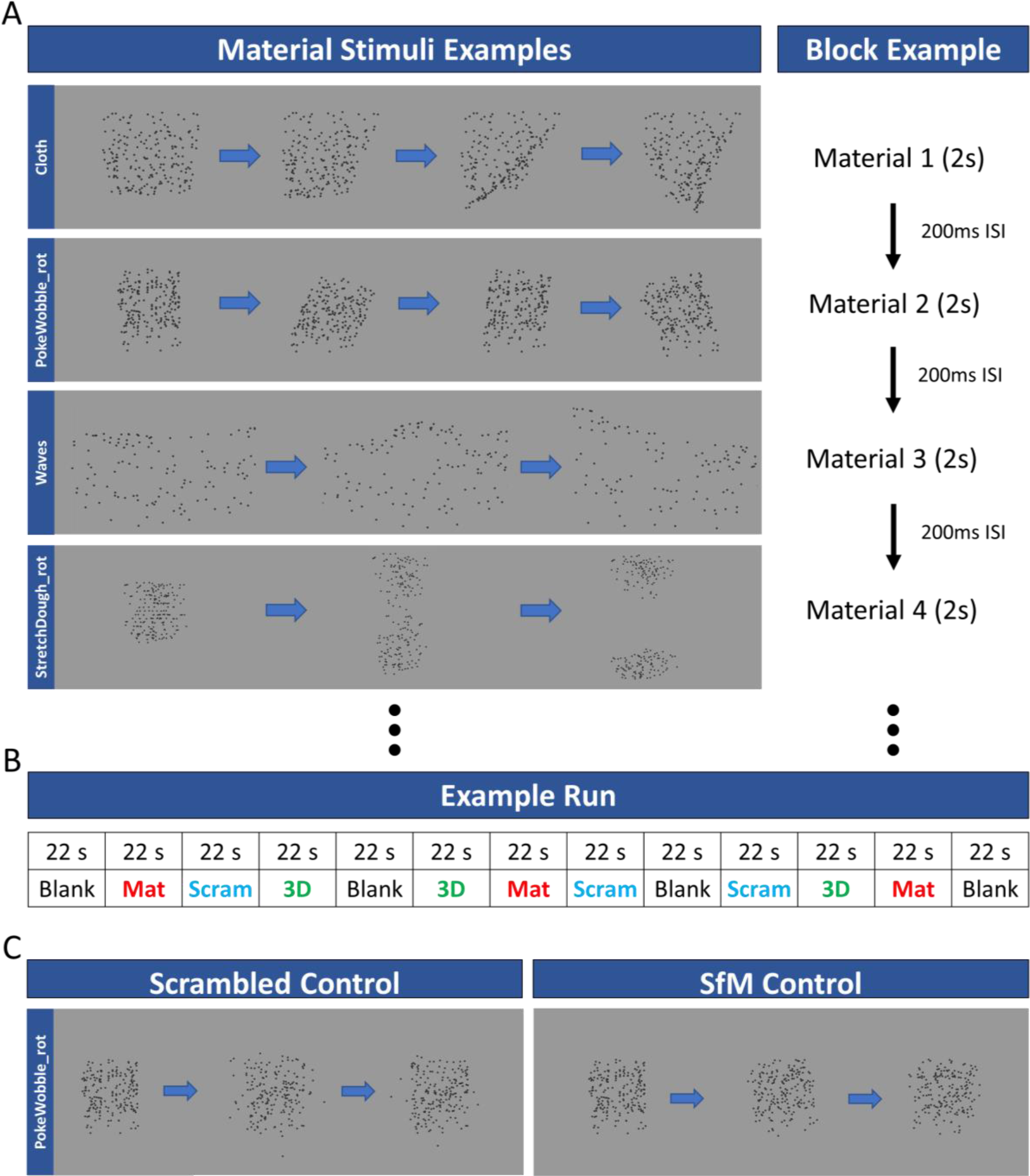
Example stimuli and block design. During the functional scans participants were presented with alternating blocks of dynamic dot materials, motion scrambled control stimuli, and structure from motion (SfM) control stimuli. On the right side of panel **A** we show selected frames of 4 of the 10 possible dynamic dot material animations that were shown in random order in a block. On the left side of the same panel the timing of presentation during a dynamic dot material block is shown. Panel **B** depicts an example run, and panel **C** shows selected frames of 1 of the 10 possible random motion control animations (left) and the SfM control stimuli (right). The timing during a block of these control conditions was identical to that of dynamic dot material blocks.

**Figure 2.**
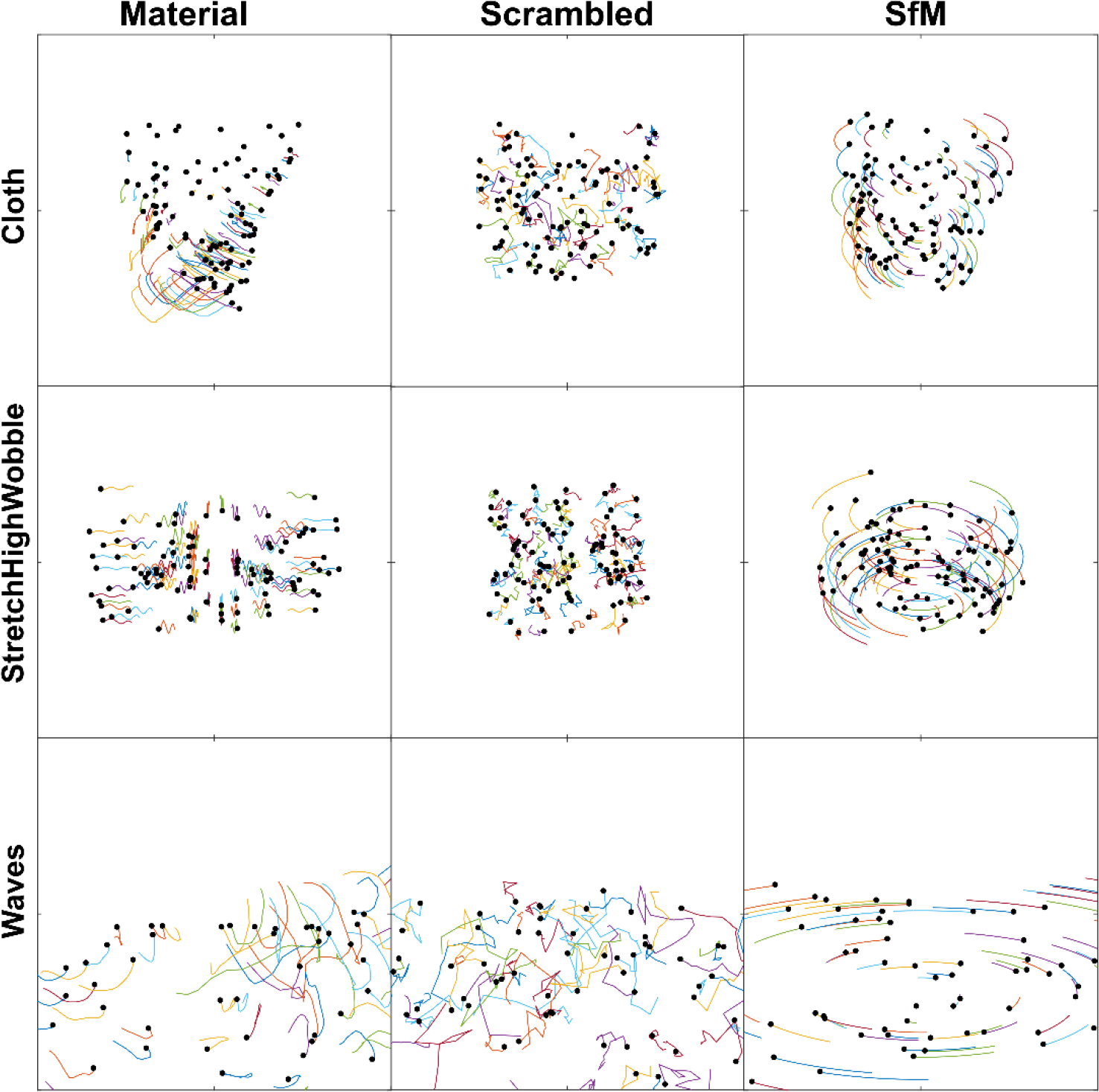
Dot Trajectories. Shown are part of the trajectories for half of the dots for three sample dynamic dot stimuli across all three conditions: material motion, velocity-matched scrambled motion control, and structure from motion (SfM) control. Each line shows the path a given dot traveled between frames 7 and 19. This also corresponds to the number of frames between reversals in the SfM condition. Trajectories for the remaining seven stimuli are shown in Supplementary Figure 1.

#### 2.2.2. Scrambled control stimuli

We wanted to find cortical regions that are sensitive to material motion over and above other kinds of motion. That is, we wanted to exclude regions that are sensitive to motion generally but do not show a preference for material motion. Therefore, we created “scrambled” control stimuli that were matched in motion energy to the dynamic dot materials (measured as velocity magnitude, see Supplementary Figures 2-4). We created a scrambled version of each material stimulus by shuffling the position of each particle on the first frame. All dots in the scrambled motion stimuli had the same velocity magnitude as the material motion; however, the direction of trajectory was rotated by a random (uniform distribution) amount between 0 and 2π every 2^nd^ frame. Without this rotation the scrambled stimuli would still look like non-rigid materials. The trajectory of a given dot was forced to remain inside the spatial range of the dynamic dot materials (in the 1^st^ frame) by forcing it to change its direction if it is outside the boundary. This was accomplished by rotating the dot trajectory to a different direction. As a consequence, acceleration between material and control stimuli were not matched (See Supplementary Figures 5-7). Example scrambled control stimuli are shown in Figure 1C, Figure 2 (middle column), and Supplementary Figure 1.

#### 2.2.3. Structure from Motion (SfM) control stimuli

We also wanted to make sure any preference for material motion within a brain region was not just a preference for global coherent 3D motion, or “object-ness”. Therefore, we created 3D structure-from-motion (SfM) control stimuli. To generate these control stimuli, we selected one frame of each material motion movie and then rotated the camera back and forth around the center of the scene. This gave the object an appearance of rotating in depth around the horizontal (or vertical axis). SfM control stimuli are shown in Figure 1C, Figure 2 (right column), and Supplementary Figure 1.

### 2.3. Stimulus display

Visual stimuli were presented on an MR-safe LCD screen placed near the rear end of the scanner bore (Cambridge Research Systems Ltd, Rochester, UK; resolution: 1920 by 1080; refresh rate 120 Hz). Participants viewed the screen through an angled mirror attached to the head-coil while lying supine inside the scanner bore. Total eye-to-screen optical distance was 140 cm, and the screen subtended a visual angle of 28 degrees horizontally at this distance. Stimuli were presented at the center of the screen and approximately subtended 15.75 by 15.75 degrees visual angle.

### 2.4. Fixation task

To ensure fixation during the scanning session, aid maintaining vigilance and wakefulness, and limit attentional effects, we asked participants to perform a demanding fixation task throughout the functional runs. In this task, participants were required to report a brief reduction in the size of the fixation cross via a button press. The cross shrank by a small amount every 3 seconds with a random (+/- up to 1.5 seconds) time jitter to make it unpredictable when the shrinking would occur.

### 2.5. MR Image Acquisition

Magnetic resonance images were collected on a 3 Tesla MRI scanner (Magnetom Prisma, Siemens AG, Erlangen, Germany) equipped with a 64-channel head coil in BION imaging center of JLU Giessen. MR sessions contained a structural run and 4-8 functional runs. Structural images were acquired using a T1-weighted 3-D anatomical sequence (sagittal MP-RAGE, Spatial resolution: 1 mm^3^ isotropic; number of slices: 176). Functional images were acquired while participants viewed the visual stimuli and were acquired with a T2*-weighted gradient-recalled echo-planar imaging (EPI) sequence (TR: 2000 ms; TE: 30 ms; spatial resolution: 3×3×3 mm^3^; number of slices: 36; slice orientation: parallel to calcarine sulcus). Each participant took part in one scanning session that lasted about an hour and a half. During functional runs participant responses were collected using an MR-safe button box.

### 2.6. Experimental Design

During the functional runs, different types of motion stimuli (material, scrambled control, SfM control) were presented in alternating blocks. The order of blocks was randomized and counterbalanced. Each block lasted 22 seconds and contained ten short clips (2 s) of animation separated by 200 ms interstimulus interval (ISI). Figure 1B depicts the experimental protocol. There were 3 repeats of each stimulus type (material, scrambled control, SfM control) per run, 4-8 runs in a session, thus 12-24 repeats of each stimulus per participant. One run lasted 286 seconds.

### 2.7. MR Data Analysis

All MR image preprocessing and further analyses were performed using BrainVoyager QX, except an initial inhomogeneity correction step on T1-weighted images, which was conducted using Freesurfer version 4 software (http://surfer.nmr.mgh.harvard.edu/). After the initial inhomogeneity correction with Freesurfer4, anatomical images were imported into BrainVoyager for further preprocessing. Preprocessing for the anatomical images included the following steps: another inhomogeneity correction using BrainVoyager (for 8 out 10 participants this led to a better white-gray matter segmentation), aligning the images in AC-PC plane and converting to Talairach space, white-gray matter segmentation. After these steps a 3D cortical mesh was created for each subject and the resulting individual meshes were morphed and aligned using cortical surface information (sulci and gyri). Finally, an average mesh was created and inflated. Preprocessing steps on functional images included slice acquisition time correction, motion correction, linear trend removal and high pass filter (temporal). The resulting functional images were coregistered with the anatomical images per participant. Functional data were spatially transformed and projected on the inflated 3D average mesh for further analyses. These analyses included whole brain random-effects surface-based group analyses using the general linear model (GLM), and region of interest (ROI) analyses.

For the whole brain analyses, we performed several separate GLM analyses (material vs. scrambled, material vs. SfM, SfM vs. scrambled). Active voxels were identified at p < 0.05 level (using BrainVoyager’s cluster correction for multiple comparisons). For the ROI analyses, we used half the data (odd runs) to define ROIs, and half the data (even runs) to extract volume time courses for each participant. ROIs were identified using fixed-effects surface-based group GLM analyses (material vs. scrambled, material vs. SfM; active voxels were identified at p < 0.05 level). For each participant and ROI, average percent signal change (from fixation baseline, using BOLD signal and middle 5 TRs from each block) was calculated for each condition (material, SfM, scrambled), and these data were subjected to a 3 (condition: material, SfM, scrambled) X 2 (hemisphere: RH, LH) repeated-measures ANOVA. Significant ANOVA results were followed up with paired t-tests using Sidak-corrected p-values for multiple comparisons (single pooled variance). We also performed sensitivity and effect size analyses for each ROI and hemisphere, which are reported in Supplementary Table 2. Sensitivity analyses were performed using G*Power (version 3.1.9.7; Faul et al., 2007), where we computed required effect size given α (0.05), Power (1-β err prob = 0.95), and sample size (n=10) for a repeated measures within-subjects ANOVA. Effect sizes were calculated as Cohen’s f (Cohen, 1988).

### 2.8. Data/code availability statement

The stimuli used in this study come from the “Dynamic dot materials database”. We provide access to this database, which contains each stimulus in the form of movies, individual movie frames, and the coordinates of each point for each frame. We also provide MR volume time course data separately for each subject, brain region, and measurement. These can be accessed through the following link: https://www.dropbox.com/sh/nakqpn022lpaptn/AACQkbsgFuVvUIZlRXnoWXrya?dl=0).

## 3. RESULTS

### 3.1 Whole Brain GLM Results

Figure 3 shows the results of the whole-brain GLM analyses. The activation map in Figure 3A shows that, overall, there are many cortical areas that have a larger BOLD response to dynamic dot materials than to scrambled motion stimuli. This widespread network of brain areas (hot colours in Figure 3A) includes dorsal and ventral visual regions in addition to multisensory, somatosensory, and premotor areas. Only early visual areas responded more strongly to the scrambled motion (cool colours in Figure 3A). This is in line with our motion energy measurements of the stimuli (Supplementary Figures 2-7) and suggests that higher activation for material versus control motion in other regions is not due to low-level differences in motion energy between the conditions (See Discussion, section 4.5: Motion energy and motion coherence).

**Figure 3.**
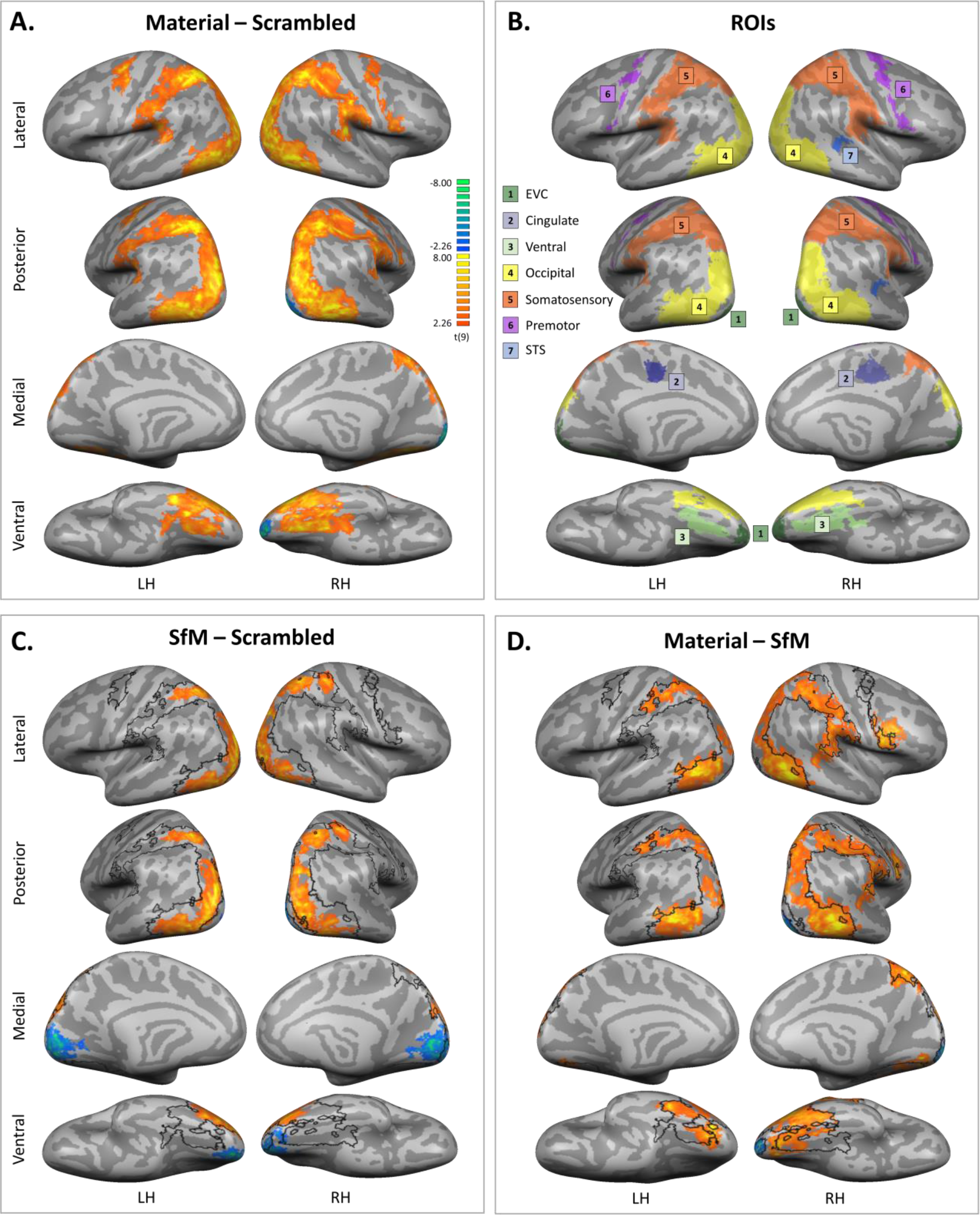
GLM group analyses. For all analyses, active voxels were identified at p < 0.05 level (cluster corrected; see section 2.7: MR Data Analyses). **A**. Results of the GLM contrasting average BOLD responses to dynamic dot materials with those to scrambled motion stimuli. Only early visual areas respond more weakly to material motion (cool colours). A large network of areas is more strongly active under the material motion condition (hot colours). The results of this contrast (material vs. scrambled) are overlaid as a black outline in (C) and (D). **B**. ROIs identified for subsequent analyses (see section 3.2: ROI Results). The ROIs largely overlap with activation maps in (A), (C), and (D). Broadly the “material motion network” encompasses ventral visual regions (3. Light green); occipito-temporal and -parietal regions (4. Yellow); dorsal visual and somatosensory/multisensory regions (5. Orange); pre-motor regions (6. Light purple); and superior temporal regions (7. Blue). **C**. Results of a GLM analysis that contrasts average BOLD responses to 3D rigid motion (SfM) with those to scrambled stimuli. The resulting activity maps suggest that cortical areas that respond strongly to material motion (black outline) are a superset of those that respond to 3D rigid motion (hot-coloured activation maps). **D**. The results of the same dynamic dot materials vs. scrambled motion contrast (black outline) together with a GLM contrast of dynamic dot materials versus 3D rigid motion (SfM; hot-coloured activation maps). Here we see that responses to dynamic dot materials were almost always stronger than those to 3D rigid motion (SfM) stimuli. See section 3.1: Whole Brain GLM Results for further details. Overall, lower visual areas tended to respond stronger to the scrambled motion stimuli. This is consistent with the literature (e.g. Murray et al. 2002).

A black outline of the activation map from Figure 3A is superimposed over the activation maps in Figure 3C and D. Figure 3C shows that cortical areas with a material “preference” (black outlines) appear to constitute a superset of those that respond to 3D rigid motion (SfM). The overlap in activation for these two contrasts (material vs. scrambled and SfM vs. scrambled) is interesting and suggests that 3D rigid motion could simply be a special type of material motion and thus activity to this type of stimuli should be contained within the general material motion network. If rigid motion is a subclass of all possible material motions then this might also explain why cortical responses to dynamic dot materials are almost always stronger than those to 3D rigid motion (Figure 3D): seeing just one type of material motion is likely to cause a relatively weaker cortical response than the rich set of motions that occur in the dynamic dot condition.

Broadly, the stronger response for material motion versus scrambled and 3D rigid motion (SfM) overlap with the results of our ROI analysis (next section). This “material motion network” encompasses ventral visual regions (light green regions labelled 3 in Figure 3B); occipito-temporal and -parietal regions (yellow regions labelled 4), dorsal visual and somatosensory/multisensory regions (though not primary somatosensory cortex; orange regions labelled 5); pre-motor regions (light purple regions labelled 6); and superior temporal regions (blue region labelled 7).

The average accuracy in the fixation task was 83% across observers, suggesting that participants followed our instructions and fixated at the center of the screen during presentation of the stimuli.

### 3.2 ROI Results

To evaluate the reliability of the results found with the whole-brain analyses, we also conducted a region of interest (ROI) analysis. Six ROIs (early visual cortex (EVC), cingulate, ventral, occipital, somatosensory, and premotor) were defined with the contrast material minus scrambled, and one ROI (superior temporal sulcus; STS) was defined with the contrast material minus SfM (active voxels were identified at p < 0.05 level). Data used to define ROIs (shown in Figure 3B) were separate to the data extracted from the ROIs for analysis (see section 2.7: MR Data Analyses). Figure 4 plots the mean percent signal change from baseline (fixation blocks) for each ROI, condition, and hemisphere. A 3 (condition: material, SfM, scrambled) X 2 (hemisphere: RH, LH) repeated-measures ANOVA revealed a main effect of hemisphere for only for the premotor ROI, F(1, 9) = 5.317, p=0.0466, where on average percent signal change was larger for the left versus the right hemisphere. There was a main effect of condition for six of the ROIs: cingulate, F(2, 18) = 6.914, p=0.0059; ventral, F(2, 18) = 19.03, p<0.0001; occipital, F(2, 18) = 59.44, p<0.0001; somatosensory, F(2, 18) = 11.43, p=0.0006; premotor, F(2, 18) = 6.886, p=0.0060; and STS^1^, F(2, 18) = 8.467, p=0.0026 (starred ROIs in Figure 4). There was a significant interaction between condition and hemisphere for two ROIs: occipital, F(2, 18) = 6.035, p=0.0099, and somatosensory F(2, 18) = 5.530, p=0.0134 (double starred ROIs in Figure 4).

**Figure 4.**
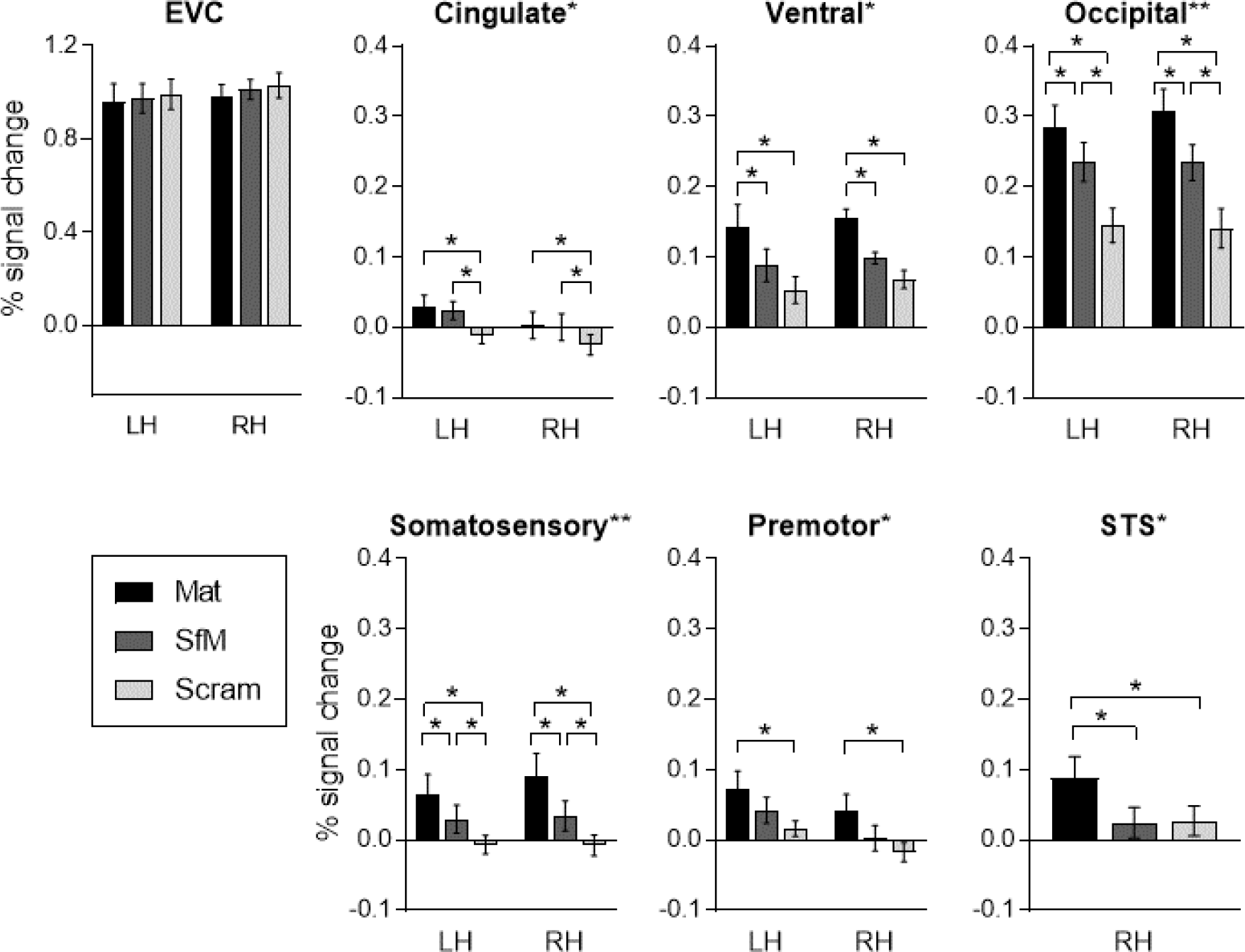
ROI analysis. Each graph shows percent signal change for each hemisphere (left hemisphere, LH; right hemisphere; RH), and each condition (Mat = material motion, SfM = shape from motion control; Scram = scrambled motion control). Starred ROI labels indicate a significant main effect of condition. Double starred labels indicate a significant interaction between condition and hemisphere. Starred contrasts indicate a significant difference between those conditions from a follow-up test (p < 0.05 level, Sidak-adjusted). See section 3.2: ROI Results for details. Error bars are one standard error of the mean and show variability between observers (see also Supplementary Figure 8 for confidence intervals associated with each contrast, and Supplementary Figures 9 and 10 for plots of each subject’s data separately).

For these two ROIs (occipital and somatosensory), follow-up tests showed significant differences between all conditions (p < 0.0001 for all tests, Sidak-adjusted p-values). That is, for both hemispheres, there was a greater BOLD response to material motion versus both control motions (SfM and scrambled), and a greater response for SfM versus scrambled motion (see Figure 4, occipital and somatosensory ROIs).

For the other ROIs where there was a main effect of condition (cingulate, ventral, premotor, and STS; single starred ROIs in Figure 4), follow-up tests revealed a larger response for material motion versus scrambled motion (p < 0.01 for all tests, Sidak-adjusted). Ventral and STS ROIs additionally showed a larger response for material motion versus SfM (p < 0.01 for all tests, Sidak-adjusted). However, the cingulate and premotor ROIs did not show a difference between these two conditions (p>0.05, Sidak-adjusted). The cingulate ROI, however, did show a greater response to SfM versus scrambled motion, suggesting that this region does not prefer material motion per se, but rather coherent motion structure.

For EVC there were no significant main effects of condition, F(2, 18) = 1.090, p=0.3574, or hemisphere, F(1, 9) = 0.3470, p=0.5703, nor a significant interaction between condition and hemisphere, F(2, 18) = 1.110, p=0.3511.

More details about the follow-up tests and effect sizes for each contrast (in the form of confidence intervals) are presented in Supplementary Figure 8. We also show that the effects reported here are extremely consistent across participants in Supplementary Figures 9 and 10.

## 4. DISCUSSION

Although motion compellingly conveys the properties of non-rigid materials such as stiffness and elasticity (e.g. Schmid & Doerschner 2018; Schmidt et al. 2017), until now the neural basis of non-rigid material motion remained unexplored. Here we investigated whether specialized dynamic dot animations portraying non-rigid material motion elicit meaningful differences in brain responses when compared to other kinds of motion using functional magnetic resonance imaging (fMRI). We found robust and widespread increased activation across the human brain in response to dynamic dot materials when compared to activation in response to other types of motion stimuli (Figures 3 and 4).

### 4.1 General activation maps in response to dynamic dot materials

Surprisingly, regions preferring dynamic dot materials included several areas in occipito-temporal and, -parietal cortices, secondary somatosensory cortex, and premotor regions. Ventral visual areas (labelled 3 in Figure 3b; light green) have previously been implicated in the processing of surfaces and textures in static scenes (Cant & Goodale, 2007; Cavina-Pratesi et al., 2010a, 2010b; Hiramatsu et al., 2011). Occipito-temporal and -parietal areas (labelled 4; yellow) include motion-, object-, face-, and place-selective areas (Grill-Spector & Mallach, 2004). Posterior parietal areas are sensitive to aspects of 3D shape, structure-from-motion, optic flow, multisensory information, and visuomotor control (Culham, et al., 2006; Erlikhman et al., 2018; Uesaki & Ashida, 2015). Recently it has been found that activity patterns in secondary somatosensory cortex (labelled 5; orange) could reliably discriminate visual properties, such as surface gloss and roughness (Sun et al., 2016b). Greater responses to biological versus scrambled point-light stimuli have been found in superior temporal areas (labelled 7; blue) and premotor areas (labelled 6; light purple) (Saygin et al., 2004). Note that the higher responses for material versus the two control motions does *not* include primary somatosensory nor primary motor cortices, but does encompass secondary somatosensory, multisensory, and premotor areas, in addition to nearly all extrastriate regions that are responsive to visual stimuli.

Discovering a network of preferentially more active areas during dynamic dot material viewing does not mean that all these areas must be involved specifically in the recognition and differentiation of materials: to find such finer-tuned responses would require further studies, and dynamic dot stimuli provide a convenient way to investigate this, as discussed next.

### 4.2 Neural mechanisms of material perception

Traditionally, material perception has been portrayed as a hierarchical processes that proceeds from the processing of image features to the perception of stimulus properties to the recognition of material identity or category (e.g. Komatsu & Goda, 2018). For example, neuroimaging studies typically investigate how particular image features (e.g. form, motion, or texture cues) might encode particular stimulus properties (e.g. 3D shape or gloss; Georgieva et al., 2008; Kam et al., 2015). A consequence of this is that stimulus properties related to material recognition are often investigated separately to one another. This separate focus on different visual properties (3D shape, texture, gloss, etc.) reflects our perceptual experience of layered visual representations and is consistent with some neuroimaging data. When we look at a scene, we interpret retinal image structure as distinct perceptual dimensions like 3D shape, surface material (gloss, transparency), colour, shading, motion, etc. (Anderson, 2011). In line with this, fMRI studies have demonstrated that different brain regions show a preference (in terms of stronger BOLD response) when attention is directed to one visual dimension (material, shape, colour, motion) over others (e.g. Cant & Goodale, 2007; Cant & Xu, 2012, 2015; 2016; Cant, Arnott, & Goodale, 2009; Caviana-Pratesi et al., 2010; Peuskens et al. 2004). This is interpreted as preferential processing of different stimulus properties in different brain regions and mirrors our perceptual experience of layered representations. If different stimulus properties are processed separately before being later integrated in a hierarchical manner (Komatsu & Goda, 2018), then to understand our multifaceted visual experience it would make sense to first study the component perceptual dimensions.

Our results challenge this view, however, as we find a preference for materials defined purely by motion properties in many areas implicated in the processing of surface material properties. For example, the ventral activity we find overlaps with posterior fusiform gyrus (pFs), fusiform gyrus (FG), collateral sulcus (CoS), lingual gyrus (LG), which have been implicated in the processing of gloss (pFs), texture (CoS, LG, FG), and material categories (FG, CoS; see Komatsu & Goda, 2018 for a review). This might suggest that these regions are not specialized for processing the image properties that depict surface properties per se, or, alternatively, that these regions are activated due to associations between material motion and surface properties (Schmid & Doerschner, 2019). We expand on this idea in section 4.3: Encoding directly perceived versus associated properties.

Our results are in line with recent neuroimaging studies that have found that regions previously associated with shape processing are also sensitive to changes in material properties (e.g. V3b/KO; Kam et al. (2015), Sun et al. (2015, 2016a); LOC, Hiramatsu et al. (2011); V4, Kim et al. (2019), as these regions overlap with occipital activation in the present study. Furthermore, our results are consistent with neuroimaging data that show crossmodal associations between visual and tactile properties. For example, visual discrimination of rough and smooth surfaces can be decoded in somatosensory cortex (Sun et al., 2016b).

The widespread activation that we find in response to quite sparse stimuli suggests that material perception is an inherently distributed process (see Schmid & Doerschner, 2019). Consistent with this idea, there is a growing body of literature suggesting that object category representations are grounded in distributed networks (e.g. Kravitz et al., 2011; 2013; Martin, 2016). Under this framework, a widespread cortical preference for dynamic dot materials is perhaps expected, given the properties of these stimuli: the stimuli are objects (LOC, e.g. Grill-Spector & Malach, 2004), they are non-rigidly moving structures (e.g. hMT+, MST, PPC, e.g. see review by Erlikhman, et al., 2018; STS, e.g. Saygin et al., 2004), they elicit a distinct tactile experience (e.g. Schmid & Doerschner, 2018; Bi et al., 2019), and such tactile experiences of material qualities are often associated with certain optical material qualities (CoS, e.g. Arnott et al., 2008; Podrebarac et al., 2014; Sun et al., 2016b; and see Komatsu & Goda, 2018 for a review).

Future work using dynamic dot materials could help reveal the underlying computations carried out by each region. For example, they could be used to investigate whether areas previously associated with materials (e.g. CoS, e.g. Cant & Goodale 2007; Cant & Xu, 2012, 2015, 2016; Gallivan et al. 2014; Eck et al., 2016; Kitada et al. 2014) are indeed specialized for processing visual (optical) properties or whether they represent material properties more generally. Our results so far suggest the latter, but further investigation, for example using a multivariate design and analysis approach, would allow one to better test this (Schmid & Doerschner, 2019). We expand on this below.

### 4.3 Encoding directly perceived versus associated properties

Materials are inherently multidimensional in that they have multiple stimulus properties that are portrayed through multiple perceptual dimensions (e.g. motion or surface optical cues), and they are inherently multimodal in that their properties can be inferred through multiple modalities (e.g. vision or touch). Over time, specific visual (or auditory, proprioceptive, or olfactory) information becomes associated with specific tactile information and vice versa. It is possible that mechanical and tactile material qualities like softness, viscosity, and roughness can be conveyed through the surface properties, 3D structure, and motion of visual objects (e.g. Baumgartner et al. 2013; Fleming, 2014, 2017; Fleming et al. 2013; Giesel & Zaidi, 2013; Ho et al. 2006; van Assen & Fleming 2016; Xiao et al. 2016). For example, a velvet cloth looks soft (optical properties), moves (visual mechanical motion) and folds (visual 3D shape) in a way that suggests that it is soft, but it also feels soft to the touch (tactile). This multidimensionality and multimodality make materials the ideal candidate to develop experimental designs that can help to understand computational architecture of the cortical representations involved in recognition. As an example, if a cortical region represents material/object category A based on visual property X but not the same category based on the visual property Y then this suggests that this region encodes the visual property X but not the category. Conversely, if this cortical region has a shared representational structure, i.e. category A is encoded through both visual properties X and Y, then it likely encodes the concept. Thus, by investigating neural responses to stimuli like the dynamic dot materials proposed here, in conjunction with stimuli defined along other visual dimensions, we may be able to tease apart the direct encoding of visual properties from the indirect activation of associated properties (Schmid & Doerschner, 2019). Note that we are referring to a generic kind of association between properties; we did not find a preference for material motion in cortical regions involved in memory or contextual associations, such as the angular gyrus, medial parietal cortex, or anterior parahippocampal cortex (Kravitz et al., 2013; Bar et al., 2008).

### 4.4 Non-rigid motion and event perception

*Dynamic dot materials* convey the non-optical properties of material qualities purely based on 2D image motion patterns. Here we introduced various types of non-rigid materials, yet also rigid materials, like breakable substances, can be rendered convincingly by the means of this technique (e.g. see Schmid & Doerschner, 2018). In their creation, these stimuli are conceptually closely related to “point light walkers” (Johannson, 1973), where the motion of small light sources affixed to different parts of limbs of biological species can elicit a very vivid impression of animacy. Bingham et al. (1995) generated point-light-like displays with retroflecting materials to capture the motion structure of animate and inanimate events, including the splashing of water or blowing of leaves in the wind. Similarly, we “attached” small dots to an otherwise invisible substance and recorded the motion of these dots while the material reacted to a force. How individual materials change their shape in response to a force strongly depends on their mechanical properties, and it is this idiosyncratic change of shape information over time that appears to convey the mechanical qualities of a material. It would be very interesting to pin-point the specific motion characteristics that elicit a particular material quality (Schmid & Doerschner, 2018; also see Bi et al. 2019), just as it has been done in the field of biological motion where researchers have tried to understand what it is that makes point-light walkers look “biological” (Chang & Troje, 2009; or Troje, 2013 for a review). The fact that the location, direction, and speed of individual dots in our stimuli can be manipulated renders investigations of research questions like these more feasible.

Such an interest intersects with general event perception, which aims to understand how we parse the continuous flow of visuo-spatial information into segments. Gunnar Johansson was a pioneer of this research and proposed a general framework of vector analysis, which relies on the decomposition of the spatiotemporal information into common and relative motion components (see Jannson et al., 1994 for an overview of Johanssons work). It is not immediately obvious what might constitute common and relative motion components in our stimuli since our shapes are rather compact, unlike a moving human or animal shape with limbs. More research is needed to understand what type of motion events observers are sensitive to (Todd, 1981). And, in pursuing this line of work, our stimuli could help to discover the perceptual boundary between non-rigid animate and inanimate objects, as well as the corresponding neural maps and mechanisms (Long, Yu, Konkle, 2018; Grill-Spector & Weiner, 2014). Moreover, since our activation maps overlap with areas also implicated in biological motion perception (e.g. STS, premotor; Saygin et al., 2004; Chang et al., 2018), our stimuli could also be suitable for teasing apart the contribution of non-rigid motion to neural activity observed while watching biological motion stimuli.

Just as all biological motion is also non-rigid motion, rigid motion is also a special case of the general event type of non-rigid motion. Our stimuli seem to ‘trigger’ this wide-spread non-rigid object motion network, with areas responding to rigid motion constituting a subset of this general network. These results are consistent with results by Bingham, et al. (1995) who found that observers can identify non-rigid types of motions as well as rigid motion events and point out, that “rigid motion is but one of a variety of types of motion” (Bingham, et al., 1995, also see Todd 1984).

Most of the recent work on event perception involves human agents or rather coarse physical interactions of objects (see Radvansky & Zacks 2011, for a review). For these kinds of scenarios it is quite intuitive to break down events into their subcomponents, like large- and small-scale events (Zacks et al. 2001). However, it is less clear what constitutes a small-scale or large-scale event in the waving motion of a cloth, or the wobbling of a jelly. This would be a very interesting topic to pursue in future work and could potentially reveal general mechanisms of event perception.

### 4.5 Motion energy and motion coherence

Our results showed that higher visual areas in the ventral and dorsal streams responded more strongly to material motion and structure from motion (SfM) over scrambled motion, and more strongly to SfM than scrambled motion, while only early visual areas (EVC) “preferred” scrambled stimuli over the other two conditions. We tested whether the preference of higher cortical areas to materials could be explained by differences in velocity of the dots in our stimuli. If we order our stimulus conditions according to the mean velocity it would yield material=scrambled=SfM; if we ordered according to the median velocity it would yield SfM>material=scrambled; and if we ordered them according to velocity range it would yield material=scrambled>SfM (Supplementary Figure 4). We can therefore rule out that differences in velocities per se drove the cortical pattern of responses we observed. In Supplementary Figure 7 we examine weather acceleration patterns can account for the measured cortical activity. Ordering the stimuli according to their acceleration profiles would yield scrambled>material>SfM. Comparing this to our fMRI measurements, we conclude that this statistic does not correspond to the observed cortical activity patterns in higher visual regions, though it might account for the preference seen for scrambled motion over SfM stimuli in early visual areas. Finding good quantitative descriptors of the motion profiles for stimuli like ours is an exciting area of research (see work in this spirit, for example in biological motion Chang & Treue 2009).

Another interesting difference between our stimulus conditions is the perception of motion coherence. Could differences in coherence explain the differences in activation we find between the conditions? This is not a trivial question, as it is not clear how we should define motion coherence: the dynamic dot materials are clearly a coherent appearance – the dots are easily grouped as belonging to the same non-rigid object – yet this coherence seems qualitatively quite different than that of rotating rigid objects (SfM stimuli). Traditional definitions involving the percentage of dots moving coherently, as for example in Shadlen et al. (1996), does not apply here. We tested several possibilities to capture coherence of our stimuli (e.g. entropy in motion direction, or divergence in optic flow), but the one that corresponds to an intuitive ordering of our stimulus classes (SfM>material>scrambled), seems to be the shape consistency across two consecutive frames. For this we resorted to a Procrustes analysis across 2 consecutive frames. The rationale was that if dots stay in their configuration (shape) across two consecutive frames it should be possible to superimpose them, i.e. match the shape of their configuration. Procrustes analysis does precisely this: it computes the linear transformation (rotation, translation scaling) between two shapes, providing an index of the error (sum of squared errors) that remains after applying this transform. We reasoned that this error should be much larger for scrambled dot stimuli, compared to the other two types of stimuli, since the configuration changes maximally between any 2 frames, and that the SfM stimuli should yield the smallest error. Supplementary Figure 11 shows the results of this analysis. Indeed, we can see that the coherence (as we defined it) in scrambled stimuli was quite low for all stimuli, and coherence of the SfM stimuli tended to be high, yet not always higher than that of the material motion stimuli. Note, that this overall pattern remains if we increase the frame distance to 4 for computing the Procrustes analysis. Overall, Supplementary Figure 11 shows that the difference in coherence between the material motion and SfM stimuli is rather small. It is possible, of course, that other measures capture a different definition of coherence (e.g. overall collinearity of motion trajectories), which we are currently investigating in a separate project.

Interestingly, Murray et al. (2002) showed that perceiving visual elements as a coherent shape increases activity in the lateral occipital complex, while at the same time reducing activity in lower visual areas, with an opposite pattern found for randomly arranged visual elements. Similarly, it is likely that the visual grouping of individually moving dots into a coherent shape is much stronger for SfM stimuli compared to randomly moving dots, potentially contributing to the stronger activation in early visual areas for scrambled stimuli. However, whether this grouping differs between material motion and SfM stimuli is not obvious, and another potential avenue to investigate in future work. Dynamic dot materials could provide a way to investigate whether areas associated with coherent motion preference (e.g. hMT+/V5 and posterior parietal cortex, e.g. Orban et al. 1999; Peuskens, 2004); show a preference for specific types of coherent motion (non-rigid vs. rigid).

## 5. CONCLUSION

Dynamic dot materials are a novel class of stimuli that convey the non-optical properties of materials purely based on 2D image motion patterns. Our results act as a proof of principle that such stimuli can be used for mapping the cortical network involved in the perception of material qualities and can act as a localizer for future studies. From a broader perspective, dynamic dot materials constitute a novel class of stimuli that holds promise for understanding general event perception, and, owing to their inherently multidimensional and multimodal nature, materials in general are a unique type of stimulus that can help neuroimaging research to advance our understanding of the computational architecture of the cortical representations involved in recognition.

## 6. ACKNOWLEDGEMENTS

This work was supported by a Sofja Kovalevskaja Award endowed by the German Federal Ministry of Education; a Marie Sklodowska-Curie Action – Innovative Training Network (MSCA-ITN/ETN) Grant, DyViTo: Dynamics in Vision and Touch – the look and feel of stuff. We thank Chris Baker for helpful comments on earlier versions of this manuscript.

## Footnotes

1 Since the STS ROI was only present in the right hemisphere, a one-way ANOVA was performed.

