## Supplementary material for "Dynamic dot displays reveal material motion network in the human brain"

### SUPPLEMENTARY FIGURES AND TABLES

| Material motion name | Ppt 1 | Ppt 2 | Ppt 3 | Ppt 4 |
|---|---|---|---|---|
| cloth | *A flag or a piece of fabric* | *Hanging (bed) sheet in the wind* | *Sheet blown by wind* | *Wind comes to a curtain* |
| Cloth_rot | *A flag* | *Hanging (bed) sheet in the wind* | *Flag blown by wind* | *Wind comes to a curtain* |
| Ripple | *Water/liquid stuff* | *Water wave* | *Rippling liquid* | *A wave went through some circles* |
| Waves | *Water / liquid stuff* | *Water wave in the ocean* | *Wave in liquid* | *[Illegible writing]* |
| pokeWobble | *A pudding* | *A jello cubic* | *Cube made out of rubber / silicone deforming* | *A ball is shaking* |
| pokeWobble_rot | *A pudding* | *A jello cubic* | *Rectangle made out of rubber / silicone deforming* | *A bottle like shape is shaking* |
| stretchBounce | *Something elastic* | *Stretched spring net* | *Stretching elastic material* | *Square shaped like a thing almost fell apart* |
| stretchWobble | *Something elastic that it breaks at the end* | *Stretched dough* | *Stretch and break of elastic material, rectangle* | *Square shaped like a thing fell apart* |
| stretchHighWobble | *Something elastic that it breaks at the end* | *Stretched jelly* | *Stretch and break of elastic material, cube* | *Square shaped like a thing fell apart* |
| stretchDough_rot | *Something elastic that it breaks in the end* | *Stretched dry dough* | *Stretch and break of elastic material, cube* | *Square shape like a thing divided into two pieces* |

***Supplementary Table 1: Four participants' perception of each dynamic dot material video.*** *Their responses are to the prompt: "Write down in a few words what you see in each movie". Note that many participants' first language was not English.*

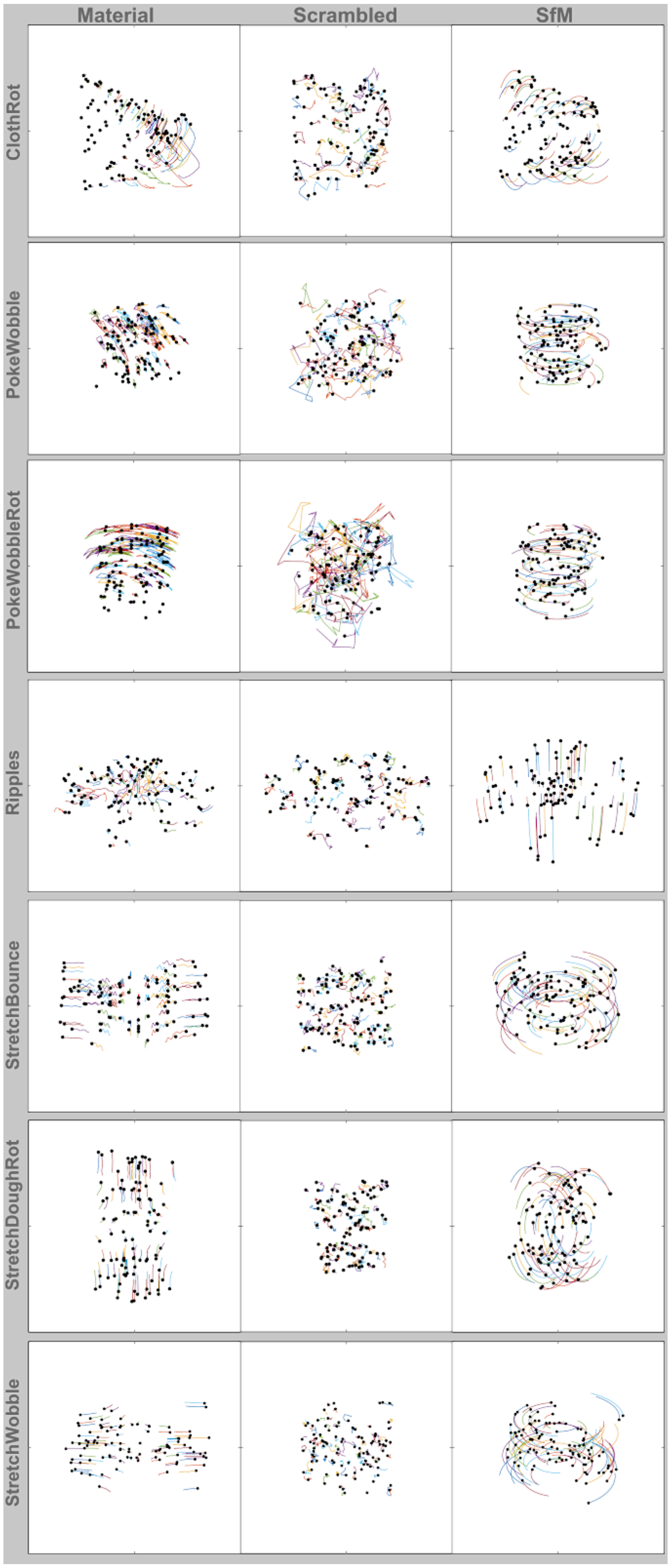


***Supplementary Figure 1. Dot Trajectories.*** *Shown are part of the trajectories for half of the dots for the remaining material stimuli across all conditions: material motion, velocity-matched scrambled motion control, and structure from motion (SfM) control. Each line shows the path a given dot traveled between frames 7 and 19. This also corresponds to the number of frames between reversals in the SfM condition. Also see Figure 2.*

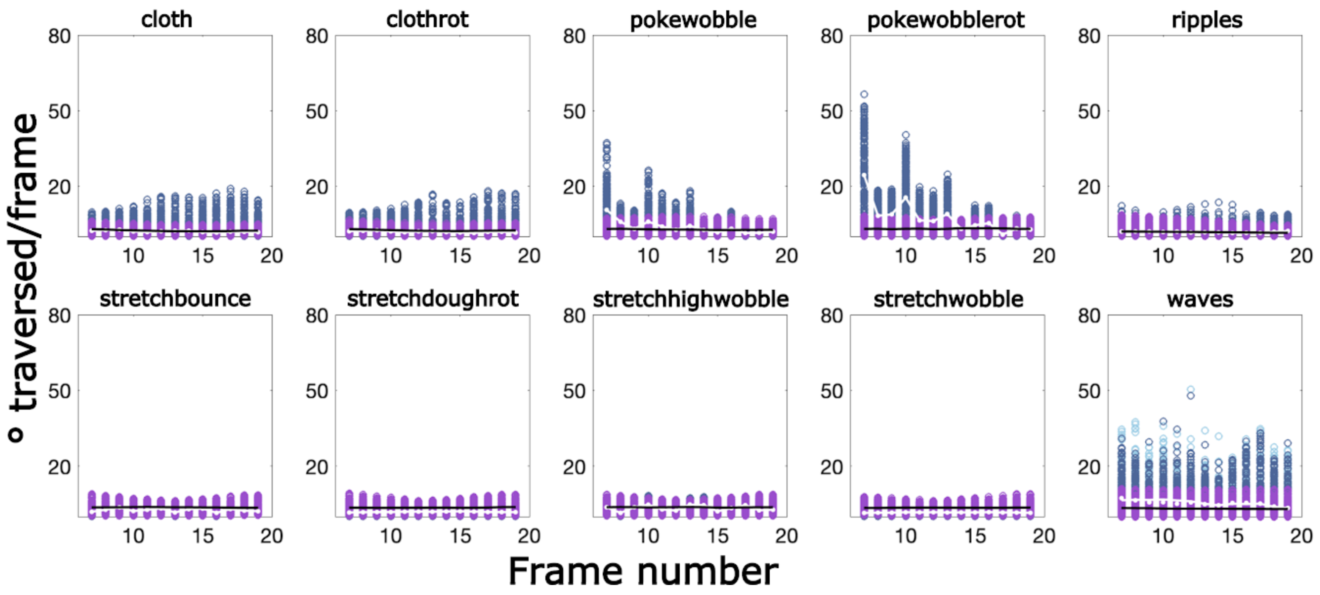


***Supplementary Figure 2: Velocities for all stimuli between frames 7 -19.*** *The Y axis shows the distance traversed by a dot in degrees visual angle between a given pair of frames. Dark blue open circles correspond to velocities of material motion stimuli, with the white line showing the corresponding velocity median, light blue open circles correspond to velocities of scrambled motion stimuli, with small white dots showing the corresponding velocity median, and purple open circles correspond to velocities of SfM stimuli, with the black line showing the corresponding velocity median. Note that for each material type the scrambled motion stimuli were velocity matched to the material motion stimuli.*

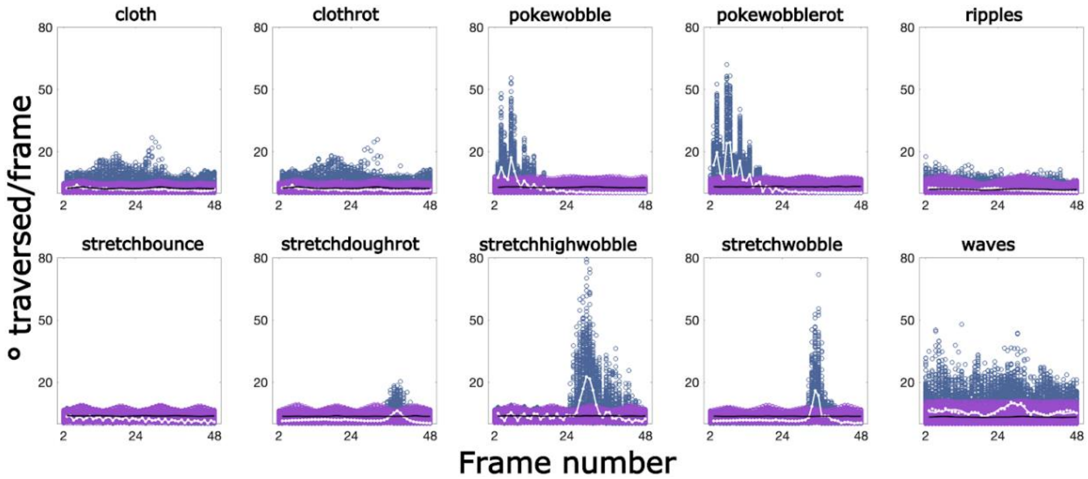


***Supplementary Figure 3: Velocities over all frames.*** *Units and symbols are the same as in Supplementary Figure 2. Note that we plot velocities from frame 2 due to an artifact in the scrambled stimuli where dots changed position (were shuffled) between 1st and 2nd frame before moving according to the profile of the matched material motion stimulus (described in the results). Swapping dot positions made no visible difference to the stimulus but did affect the coordinates that we supply with material motion database.*

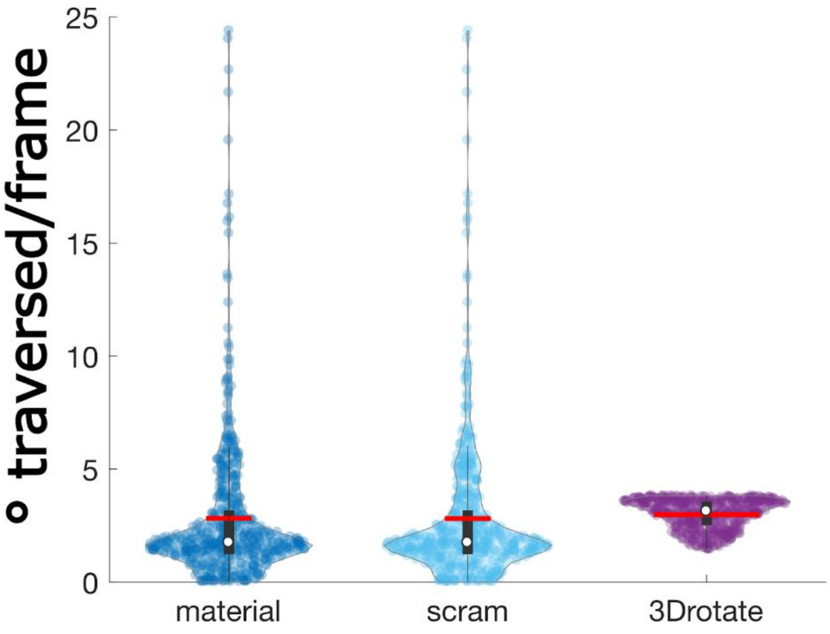


***Supplementary Figure 4: Velocities.*** *Violin plots showing the distribution of velocities for each condition across all material types. Transparent dots are the actual velocity samples. Red lines indicate the mean, white dots the median. This depiction captures some interesting motion characteristics of each stimulus type (material, scrambled, SfM). The mean of all three stimulus types (red lines) are rather similar; however the distributions of dynamic dots (dark blue) and scrambled dots (light blue) are skewed. Therefore the median (white dot) is much more informative about the central tendency of velocity: here we can see that the median velocity of SfM dots was somewhat higher than for the other two stimulus categories. This plot also shows that the range of velocities differs between stimulus types, with SfM stimuli exhibiting a much smaller range of velocities (black bars denote interquartile range; thin, central lines the whiskers, dots beyond the whiskers are considered outliers). We can see that velocities in the SfM are distributed much more homogeneously than in the other two stimulus conditions.*

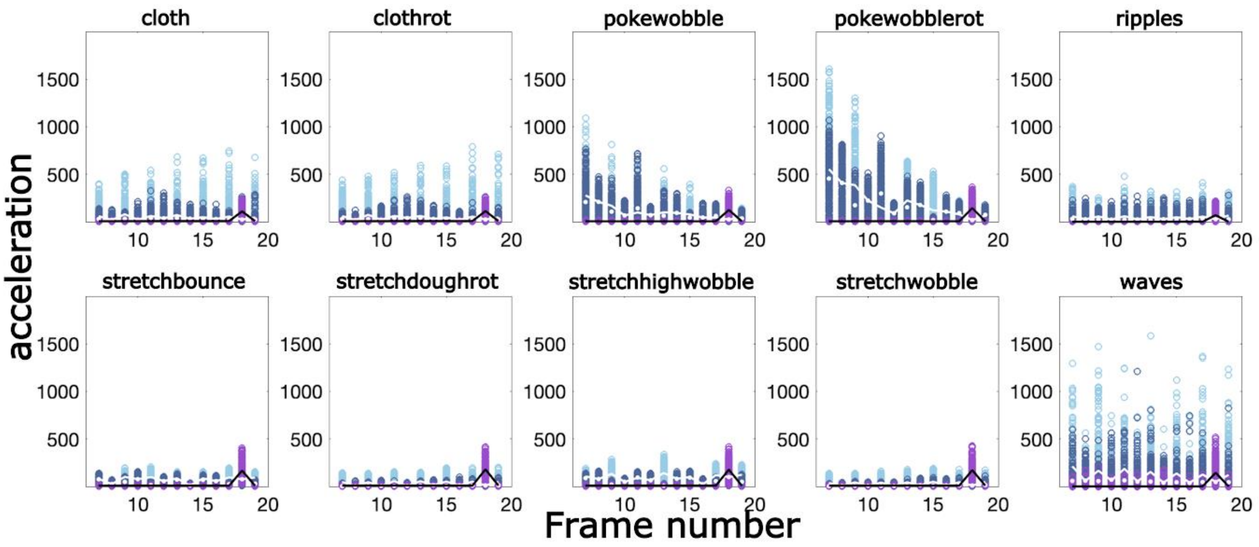


***Supplementary Figure 5: Acceleration distribution frames 7-19.*** *Acceleration is computed as the difference in velocity between two consecutive frames. Symbols and colors are as in Supplementary Figures 2-4. Acceleration was not matched between material motion (dark blue circles, white line) and scrambled motion (light blue circles, white points) conditions, since the scrambled dots changed their trajectory randomly on every frame. Supplementary Figure 6 below shows acceleration distributions for each material type, across all conditions and across all frames.*

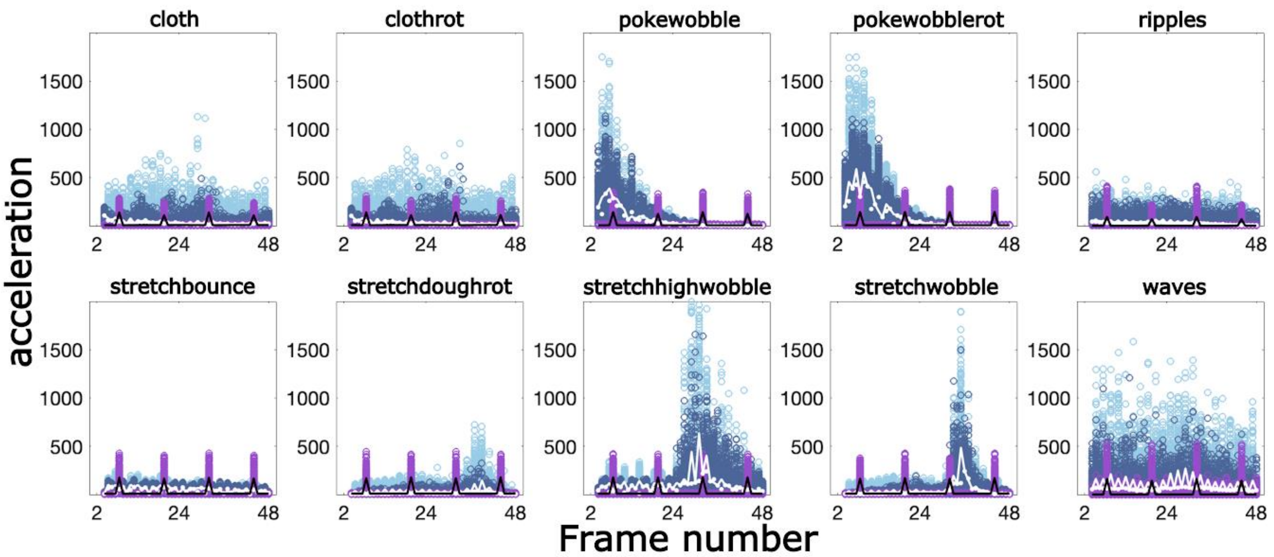


***Supplementary Figure 6: Acceleration distribution across all frames.*** *Symbols and colors are as in Supplementary Figures 2-5. Overall we can see accelerations differ across frames, especially for material motion and scrambled motion stimuli. Spikes in the acceleration profile of SfM stimuli (purple circles, black line (median)) correspond to the points when the rotation of the SfM stimuli reversed their direction. This occurred every 12 frames (frame: 7, 19, 31,43), i.e. one reversal every 2 seconds.*

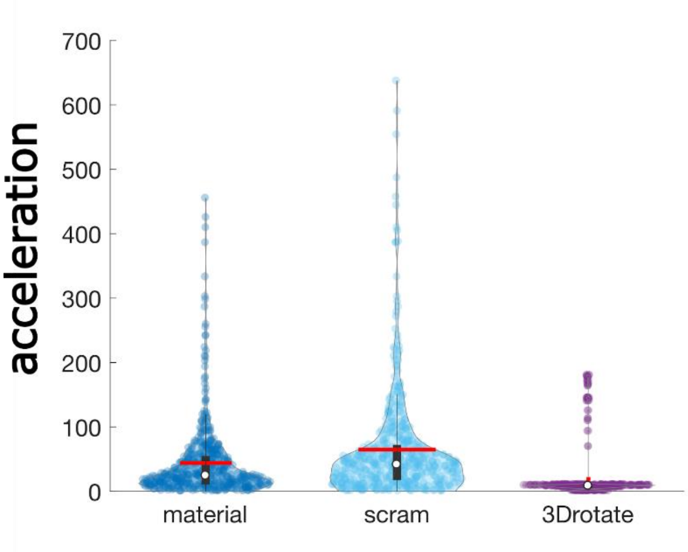


***Supplementary Figure 7: Acceleration distributions.*** *Violin plots intuitively summarize the distribution of accelerations shown in Supplementary Figure 6 above: on average (red lines) accelerations tended to be higher for material motion and scrambled motion stimuli compared to SfM stimuli. This same ranking was true for the median and the range of accelerations. Ordering the stimuli according to their acceleration profiles would yield: scrambled>material>SfM. Comparing this to our fMRI measurements we conclude that also this statistic does not correspond to the observed cortical activity patterns. Overall one can see that such summary statistics capture the actual velocity and acceleration patterns only to some extent.*

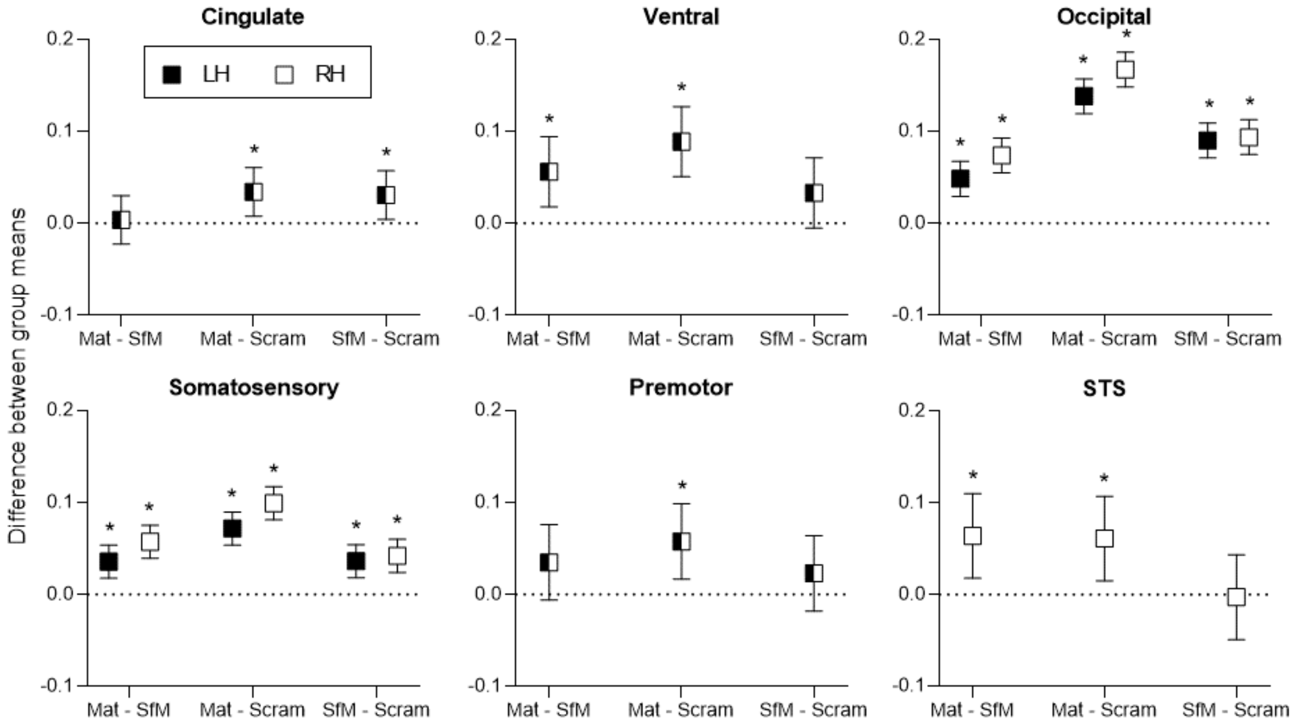


***Supplementary Figure 8: 95% confidence intervals*** *for the follow-up tests performed on the ROI data (Sidak-adjusted; see section 3.2: ROI Results). Corresponding statistical tests are as follows (for all tests df = 18):* ***Cingulate:*** *Mat-SfM (t=0.3528, p=0.9800), Mat-Scram (t=3.382, p=0.0099), SfM-Scram (t=3.030, p=0.0215);* ***Ventral:*** *Mat-SfM (t=3.845, p=0.0036), Mat-Scram (t=6.100, p<0.0001), SfM-Scram (t=2.256, p=0.1063);* ***Occipital LH:*** *Mat-SfM (t= 7.496, p<0.0001), Mat-Scram (t=21.48, p<0.0001), SfM-Scram (t=13.98, p<0.0001);* ***Occipital RH:*** *Mat-SfM (t=11.44, p<0.0001), Mat-Scram (t=25.99, p<0.0001), SfM-Scram (t=14.55 13.98, p<0.0001);* ***Somatosensory LH:*** *Mat-SfM (t=5.818, p<0.0001), Mat-Scram (t=11.72, p<0.0001), SfM-Scram (t=5.899, p<0.0001);* ***Somatosensory RH:*** *Mat-SfM (t=9.332, p<0.0001), Mat-Scram (t=16.18, p<0.0001), SfM-Scram (t=6.850, p<0.0001);* ***Premotor:*** *Mat-SfM (t=2.223, p=0.1133), Mat-Scram (t=3.685, p=0.0051), SfM-Scram (t=1.462, p=0.4092);* ***STS:*** *Mat-SfM (t=3.642, p=0.0056), Mat-Scram (t=3.480, p=0.0080), SfM-Scram (t=0.1614, p=0.9980).*

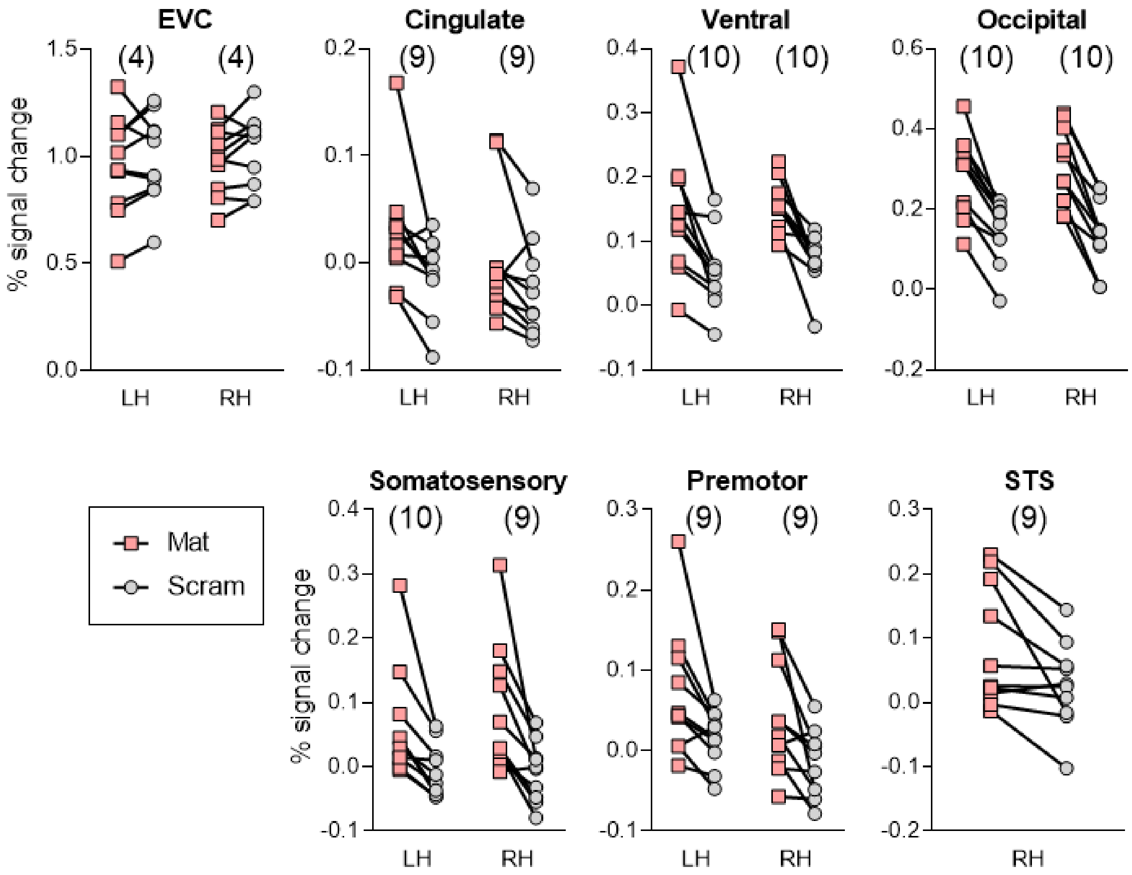


***Supplementary Figure 9: Percent signal change plotted individually for each subject**, for material motion (Mat) and scrambled motion (Scram) conditions. The numbers in parentheses show the number of participants whose BOLD response was greater for material motion versus scrambled motion for that ROI and hemisphere. Note that the axes differ for each plot for better visualization.*

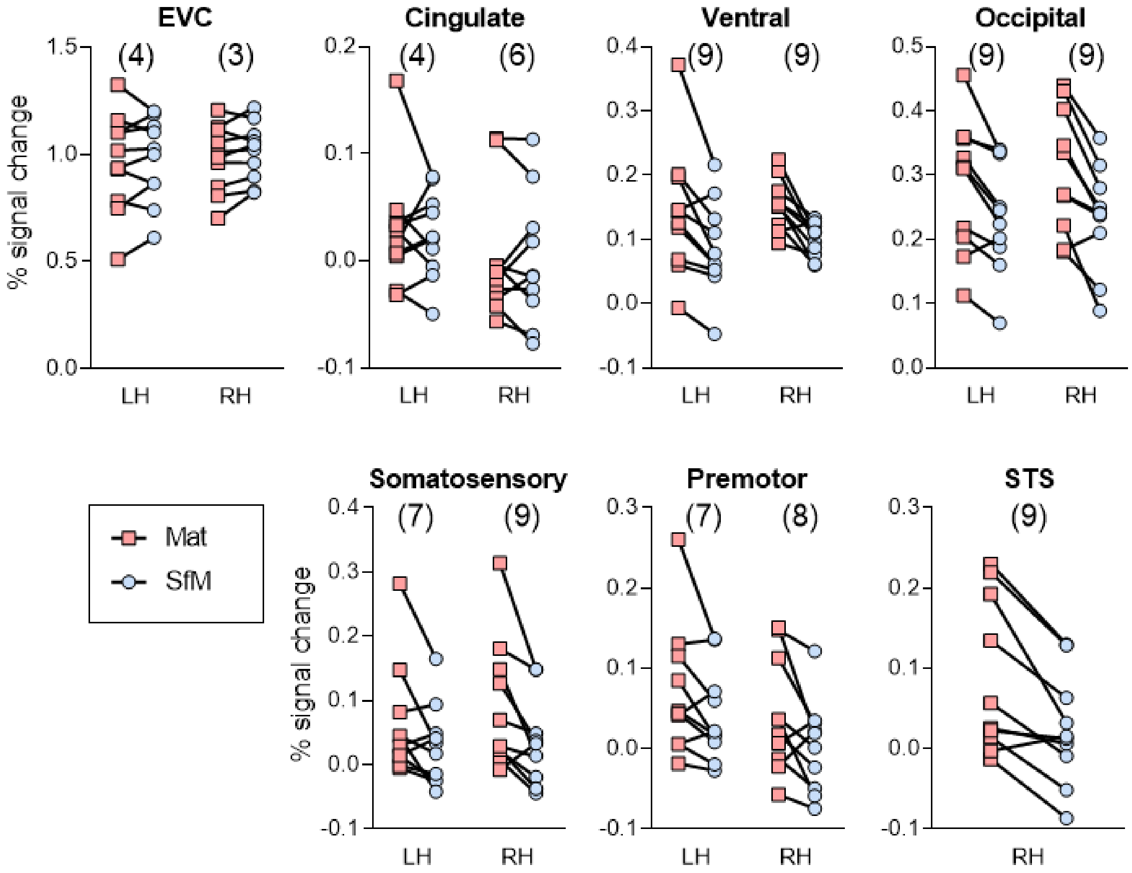


***Supplementary Figure 10: Percent signal change plotted individually for each subject**, for material motion (Mat) and rigid structure from motion (SfM) conditions. The numbers in parentheses show the number of participants whose BOLD response was greater for material motion versus SfM for that ROI and hemisphere. Note that the axes differ for each plot for better visualization.*

| ROI | Hemi-sphere | $\rho_{av.}$ | Sensitivity (Effect size f) | Sensitivity effect size mag | Cohen's f | Detected effect size mag |
|---|---|---|---|---|---|---|
| EVC | LH | 0.91 | 0.23 | small | 0.07 | small |
| | RH | 0.90 | 0.25 | small | 0.15 | small |
| Cingulate | LH | 0.69 | 0.44 | large | 0.49 | large |
| | RH | 0.86 | 0.29 | med | 0.27 | med |
| Ventral | LH | 0.84 | 0.31 | med | 0.56 | large |
| | RH | 0.43 | 0.59 | large | 1.22 | large |
| Occipital | LH | 0.89 | 0.26 | med | 0.78 | large |
| | RH | 0.87 | 0.28 | med | 0.94 | large |
| Somat. | LH | 0.82 | 0.33 | med | 0.53 | large |
| | RH | 0.72 | 0.41 | large | 0.66 | large |
| Premotor | LH | 0.79 | 0.36 | med | 0.45 | large |
| | RH | 0.63 | 0.48 | large | 0.52 | large |
| STS | RH | 0.79 | 0.36 | med | 0.46 | large |

***Supplementary Table 2: Sensitivity analysis and effect sizes for each ROI.*** *This table presents a comparison of the minimum detectable effect size given our sample size and correlation among measures, alongside the effect sizes obtained in the experiment. "$\rho_{av}$" is the average correlation among repeated measures. "Sensitivity (Effect size f)" is the minimum detectable effect size given our given sample size and correlation among measures ($\rho_{av}$). "Sensitivity effect size mag" is an evaluation of the magnitude of the minimum detectable effect size. "Cohen's f" is the effect size that was detected in the experiment. "Detected effect size mag" is the magnitude of the effect size that was detected. Evaluation of effect sizes are based on Cohen's criteria: values of 0.10, 0.25, and 0.40 represent small, medium, and large effect sizes, respectively.*

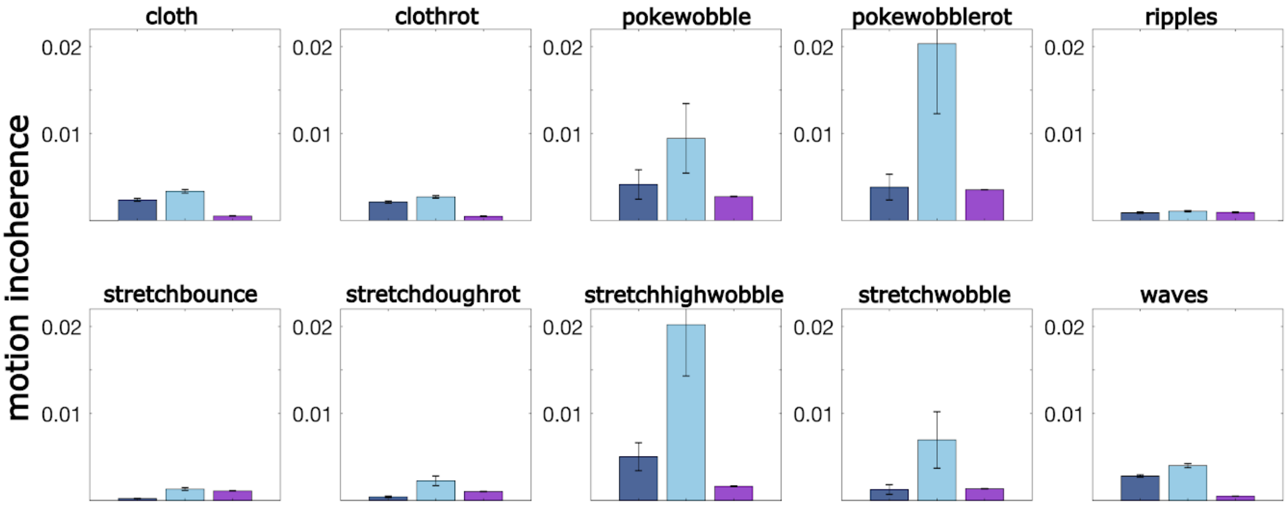


***Supplementary Figure 11: Motion incoherence.*** *Despite roughly matched velocity statistics our stimuli differed in appearance, such as coherence. In an attempt to quantitatively capture this qualitative difference in appearance we defined coherence as the degree to which the configurational shape of the distribution dots in two consecutive frames can be matched using a linear transform (rotate translate, scale). Using a procrustes analysis we computed for each pair of frames this transform and the remaining error. The average of this error across all frame pairs is plotted here and we call it motion incoherence. Color codes are as in Supplementary Figures 2-7. As expected motion incoherence was largest for scrambled stimuli. Incoherence of material motion stimuli tended to be somewhat higher than that of SfM stimuli, although this difference was small and sometimes reversed for some material types (pokewobblerot, ripples, stretchbounce, or stretchwobble).*
